# Differences in brain activity during sentence repetition in people who stutter: a combined analysis of four fMRI studies

**DOI:** 10.1101/2025.07.09.663990

**Authors:** Birtan Demirel, Jennifer Chesters, Emily Connally, Patricia Gough, David Ward, Peter Howell, Kate E. Watkins

**Affiliations:** Department of Experimental Psychology, University of Oxford, Oxford, United Kingdom, OX2 6GG; School of Psychology, University College Dublin, Dublin, Ireland, D04 V1W8; School of Psychology & Clinical Language Sciences, University of Reading, Reading, United Kingdom, RG6 6AL; Experimental Psychology, Psychology & Language Sciences, University College London, London, United Kingdom, WC1E 6BT

**Keywords:** stuttering, speech disorder, brain imaging, fMRI, people who stutter

## Abstract

Our understanding of the neural correlates of developmental stuttering benefits from the use of functional MRI (fMRI) during speech production. Despite two decades of research, however, we have reached little consensus. In the current study, we analysed pooled fMRI data from four different studies that used the same sentence reading task and methodological approach. The combined sample included 56 adolescents and adults who stutter and 53 demographically matched typically fluent controls. A sparse-sampling design was used in each study, in which participants spoke during the silent period between measurements of brain activity. Sentence reading evoked activity in both groups across frontal and temporal regions bilaterally. At statistical thresholds corrected for family-wise error, there were no significant group differences. An uncorrected threshold was applied to explore group differences in areas previously identified in earlier fMRI studies on stuttering. People who stutter (PWS) showed greater activity compared with controls in right frontal pole, right anterior insula extending to frontal operculum, left planum temporale, and midbrain, at the level of red nucleus. In contrast, PWS showed lower activity in left superior frontal sulcus, subgenual medial prefrontal cortex, right anterior temporal lobe, and portions of inferior parietal lobe bilaterally including the angular gyrus on the left. Despite pooling data across multiple studies to achieve a relatively large sample, group differences in regions involved in speech-motor control only emerged at an uncorrected voxel-wise threshold. Some of these findings align with previous fMRI studies, such as increased activity in the right anterior insular cortex.

## Introduction

Stuttering is a complex neurodevelopmental disorder that affects about 1% of the adult population worldwide. It is characterised by disruptions in the flow of speech, with symptoms appearing in early childhood, usually between the ages of two and four years (Yairi & Ambrose, 2013). Recent neuroimaging studies have significantly improved our understanding of the brain regions involved in stuttering, showing both structural and functional differences in areas responsible for speech motor skills, sensorimotor integration, error monitoring, cognitive functions, and emotional regulation (for a comprehensive review: Neef & Chang, 2024). However, there is still no definitive consensus on the neural correlates of stuttering.

Meta-analytical approaches applied to fMRI and positron emission tomography (PET) data have deepened our knowledge of functional differences between PWS and typically fluent speakers. By combining results from multiple studies, three different meta-analyses attempted to identify the neural correlates of stuttering during speech tasks (Belyk et al., 2015; Brown et al., 2005; Budde et al., 2014). The first meta-analysis, encompassing eight PET and fMRI scans, revealed three key ‘neural signatures’ of stuttering: overactivity of right anterior insula/frontal operculum, underactivity of the auditory cortex, and overactivity of cerebellar vermis (Brown et al., 2005). Reduced activity in the auditory cortex is thought to result from mismatches between auditory feedback and the efference copy of intended speech motor commands in the sensorimotor integration system (Civier et al., 2010; Max et al., 2004). Resting-state and intervention studies suggested a potential compensatory role for the cerebellum, with stronger cerebellar-frontal connectivity associated with milder stuttering (Sitek et al., 2016), and fluency therapy linked with increased cerebellar activity (De Nil et al., 2001; Kell et al., 2018).

Following these initial findings, subsequent meta-analyses have produced mixed results in their trait-related analysis, in which they compared brain activity during fluent speech between PWS and fluent speakers. One meta-analysis failed to replicate the cerebellar vermis finding but confirmed increased activity in the right insula and decreased activity in the left temporal lobe in PWS (Budde et al., 2014). It also reported novel findings of increased activity in supplementary motor area (SMA), which plays a role in movement initiation, and decreased activity in the left red nucleus. Belyk and colleagues conducted a similar meta-analysis in 2015, which was corrected subsequently for implementation errors that had affected statistical thresholds due to a software glitch. Their updated analysis retained few robust effects from earlier studies, failing to replicate previous cerebellum and temporal lobe findings but identifying increased activity in the right Rolandic operculum and decreased activity in the left orofacial pre/primary motor cortex, which are key brain regions involved in articulation (Belyk et al., 2017).

Another set of critical structures implicated in stuttering are the basal ganglia, which are thought to be involved in the selection, initiation, and termination of speech motor commands, while inhibiting competing ones (Alm, 2004; Craig-McQuaide et al., 2014, Chang & Guenther, 2020). This connection between stuttering and the basal ganglia emerged after an early imaging study identified elevated dopamine levels in the striatum of three PWS compared with six typically fluent controls (Wu et al., 1997). Soon afterwards, an fMRI study found that baseline activity in the caudate nucleus bilaterally correlated positively with stuttering severity during fluent speech in the scanner, and that this correlation decreased after fluency-shaping therapy (Giraud et al., 2008). Furthermore, another fMRI study reported increased activity in a large subcortical area at the top of the brainstem, encompassing several structures, including the substantia nigra, pedunculopontine nucleus, red nucleus, and subthalamic nucleus (Watkins et al., 2008). These individual fMRI studies suggested that some functional differences at the subcortical level were not captured by meta-analyses. Moreover, recent lesion network analysis revealed that neurogenic stuttering and persistent developmental stuttering share a common neural network, focused on the left-sided posteroventral putamen (Theys et al., 2024). Evidence for the causal role of the basal ganglia in stuttering comes from findings that deep brain stimulation in the globus pallidus, pars interna (GPi) in Parkinson’s patients can induce stuttering (Nebel et al., 2009; Rusz et al., 2018). Levodopa treatment, which increases dopamine levels, can also exacerbate stuttering (Louis et al., 2001; Tykalová et al., 2015) possibly by making it difficult to select the correct motor program over incorrect alternatives (Chang & Guenther, 2020).

Despite growing evidence from task-based, resting-state, and intervention studies, a clear consensus on the functional neural correlates of stuttering has yet to emerge, as findings remain inconsistent in terms of brain regions, across imaging modalities, and methodological approaches. Contributing factors to this variability include experimental tasks (e.g., sentence reading, picture description), imaging techniques (e.g., PET, fMRI), methodological approaches (e.g., sparse vs. continuous sampling), and the use of fluency enhancement techniques (e.g., metronome pacing, choral speech). To reduce such variability, we conducted a combined analysis of four fMRI studies that used the same speech task and methodological approach, collected over time by the same research group. This approach minimises the confounding effects of individual study design differences and leverages a larger dataset to enhance the statistical power of our findings.

We pooled data from four different studies, three of which have been published (Watkins et al., 2008; Connally et al., 2018; Chesters et al., 2021, preprint) (see Table 1 for scan details). This allowed us to apply a uniform processing pipeline and stringent whole-brain correction for multiple comparisons to a relatively large sample. This provided a more reliable test of previously reported activity differences, addressing prior differences in sample size and analysis thresholds. The study of Watkins and colleagues (2008), revealed overactivity in subcortical regions such as the midbrain, potentially related to basal ganglia disfunction (since the areas encompassed the substantia nigra, see above), and underactivity in cortical motor and premotor areas associated with articulation and speech production. Connally et al. (2018) focused on differentiating state (e.g., stuttering events) and trait effects (e.g., persistent differences in brain activity related to stuttering) in stuttering using fMRI. Their findings identified reduced activity in auditory and motor regions as general traits of stuttering, while disfluent states showed greater activity in inferior frontal and premotor areas, extending into the frontal operculum bilaterally. Lastly, Chesters et al. (2021, preprint) examined neural changes associated with a five-day intervention combining anodal transcranial direct current stimulation over the left inferior frontal cortex with fluency-enhancing techniques. At the pre-intervention baseline, PWS showed lower activity compared with controls in several regions, including the right subcentral gyrus, medial parietal cortex extending to paracentral lobule, and portions of the anterior cerebellum; while there were no regions where PWS showed higher activity than controls at baseline. Therefore, based on previous meta-analyses and the individual studies we pooled, we hypothesised that PWS would exhibit differences in neural activity compared with controls. Specifically, we anticipated overactivity in regions such as the right anterior insular cortex, and cerebellar vermis, alongside underactivity in auditory cortex, and motor and premotor regions.

**Table 1:** Acquisition details for each study. The table summarises the scanning parameters used in the four studies included in the analysis. TE = Echo time (ms); TR = Repetition time (s); TA = Acquisition time (s); acquisition is delayed in sparse sampling designs to allow a silent interval between acquisitions resulting in a longer TR (= delay + TA); PWS: People who stutter; CON: Controls.

| Study | Baseline | PWS | CON | Scanner | TE (ms) | TR/TA (s) | Slices (N) | Thickness (mm) | In-plane (mm) |
| --- | --- | --- | --- | --- | --- | --- | --- | --- | --- |
| Watkins et al., 2008 | Row Xs | 10 | 10 | 3T Varian-Siemens | 30 | 10 / 3 | 32 | 4 | 4 x 4 |
| Unpublished data | Row Xs | 7 | 10 | 3T Siemens Trio | 30 | 10 / 3 | 50 | 3 | 3 x 3 |
| Chesters et al., 2021 (preprint) | Farsi script | 23 | 15 | 3T Siemens Trio | 30 | 9 / 2 | 38 | 3.5 | 3 x 3 |
| Connally et al., 2018 | Fixation | 16 | 18 | 3T Siemens Trio | 30 | 9 / 2 | 32 | 4 | 3 x 3 |

## Methods

### Participants

fMRI data were collected from four different cohorts over the past two decades, including both adolescents and adults diagnosed with stuttering, and matched control groups consisting of typically fluent speakers. The data were gathered during overt sentence reading tasks (see Demirel et al., 2024 for language laterality analysis of the same dataset). After applying exclusion criteria for excessive movement during scanning (detailed below), data in 5 PWS were excluded from the analysis, leaving useable datasets from 56 PWS (10 females, 46 males; 7 left-handed; mean age = 28 years; age range = 14-54 years) and 53 control participants (13 females, 40 males; 2 left-handed; mean age = 28 years; age range = 14-53 years). Chi-squared tests indicated no significant differences between the groups in terms of handedness distribution (chi-squared = 2.73, p = 0.10) or gender distribution (chi-squared = 0.73, p = 0.39). Within the PWS group, stuttering severity, as measured by the Stuttering Severity Instrument (SSI) (Riley, 1994), ranged from very mild to very severe (range 9-46, median 26). The classifications of stuttering severity among the PWS participants were as follows: very mild (n = 7), mild (n = 10), moderate (n = 23), severe (n = 11), and very severe (n = 5).

### Speech task

The current analysis focused on the sentence reading task, as it constituted the largest dataset we could assemble from the four individual studies (Table 1). Sentences were displayed in white font against a black background. In one study (16 PWS, 18 CON) (Connally et al., 2018), a scene relevant to the sentence was also presented concurrently. Participants were required to read the sentences aloud during the quiet intervals between scanning measurements and were able to listen to their own voice through headphones. Low-level baseline conditions were established slightly differently in each study: a series of letter Xs substituting for all characters in a sentence (17 PWS, 20 CON) (Watkins et al., 2008), sentences composed in the Farsi alphabet, which were not decipherable by the participants (23 PWS, 15 CON) (Chesters et al., preprint), and a fixation cross (16 PWS, 18 CON) (Connally et al., 2018). These baseline conditions created a contrast between the visual stimulation of symbols with the overt reading of meaningful sentences, allowing us to identify task-evoked activation patterns relevant to visual word recognition, semantic and syntactic processing, phonological assembly, articulation, and monitoring of self-produced speech.

### FMRI data analysis

FMRI data were analysed at the single-subject and group levels using FEAT (fMRI Expert Analysis Tool) from the FMRIB Software Library (FSL; Woolrich et al., 2001). Echo-planar imaging (EPI) sequences went through several pre-processing steps. EPIs were corrected for head motion using FSL’s MCFLIRT tool (Jenkinson et al., 2002), and spatial smoothing was applied using a Gaussian kernel with an 8-mm full-width at half-maximum. Exclusion criteria were defined such that any dataset displaying an absolute mean head displacement greater than 4 mm (exceeding the largest voxel dimension), or scans flagged by MCFLIRT for excessive motion during the session, were excluded. Following these criteria, scans from five participants within the PWS group were omitted from further analysis; no datasets from control participants were excluded.

A high-pass filter with a cut-off of 90 seconds was applied to remove low-frequency noise. Where available, field map-based distortion correction was performed to correct for magnetic field inhomogeneities. Subsequently, each participant’s functional images were co-registered to their individual high-resolution T1-weighted structural images via boundary-based registration.

For further standardisation across the study cohort, these individual anatomical images were then registered to the standard MNI-152 brain template using the non-linear registration algorithm FNIRT, which is also part of FSL. To account for residual effects of head motion, six parameters obtained during the motion correction phase were included in the subsequent analyses as covariates of no interest.

The study employed a sparse temporal sampling technique (Hall et al., 1999) to minimise scanner noise interference during speech production. Participants were instructed to speak only during silent intervals between scan acquisitions. Sparse-sampling acquisitions have been shown to outperform continuous imaging in capturing auditory responses in the temporal lobe cortex (Blackman & Hall, 2011; Gaab et al., 2007). Data acquisition is timed to capture the peak of the haemodynamic response to an event, which typically occurs 4 - 6 seconds later (Glover, 1999).

### Statistical analysis

Task-related blood-oxygen-level-dependent (BOLD) activity was quantified using the general linear model in FSL’s FEAT. For each participant, contrasts of interest were defined to compare the mean BOLD signal during overt sentence reading against the baseline condition. We used FMRIB’s Local Analysis of Mixed Effects stage 1 (Woolrich et al., 2001) for the group-level analyses, with the study origin included as a covariate to account for potential variability between the different datasets pooled. Statistical inference involved using a cluster-forming threshold of Z > 3.1 and an extent threshold of p < 0.05, with family-wise error correction for multiple comparisons. We examined group differences (PWS vs. CON) in task-related activity (sentence reading) relative to baseline.

When no significant differences in activity between groups were observed at this corrected threshold, we employed an exploratory approach using an uncorrected threshold of p < 0.05 (Z > 2.3), with the condition that clusters must include at least 30 voxels (k > 30). To maximise power, we included all speaking trials in this analysis. Had we found robust group differences, we planned to explore how these changed once stuttered epochs were excluded. Given the lack of differences, however, this planned analysis was not done (see discussion section for a more detailed discussion).

## Results

### Group averages

First, we analysed the BOLD activity for each group in comparison to the baseline conditions. Results indicated a similar pattern of broad activation in both groups across brain regions typically associated with speech and language production (Fig. 1). Specifically, extensive activity was observed bilaterally in sensorimotor cortex, ventral premotor cortex, inferior frontal gyrus (though more extensive on the left than the right) and pre-supplementary motor area, extending to the anterior cingulate cortex. Additional significant activity was noted in superior temporal lobe, thalamus, ventral occipito-temporal cortex, and lobule VI of the cerebellum.

**Figure 1:**
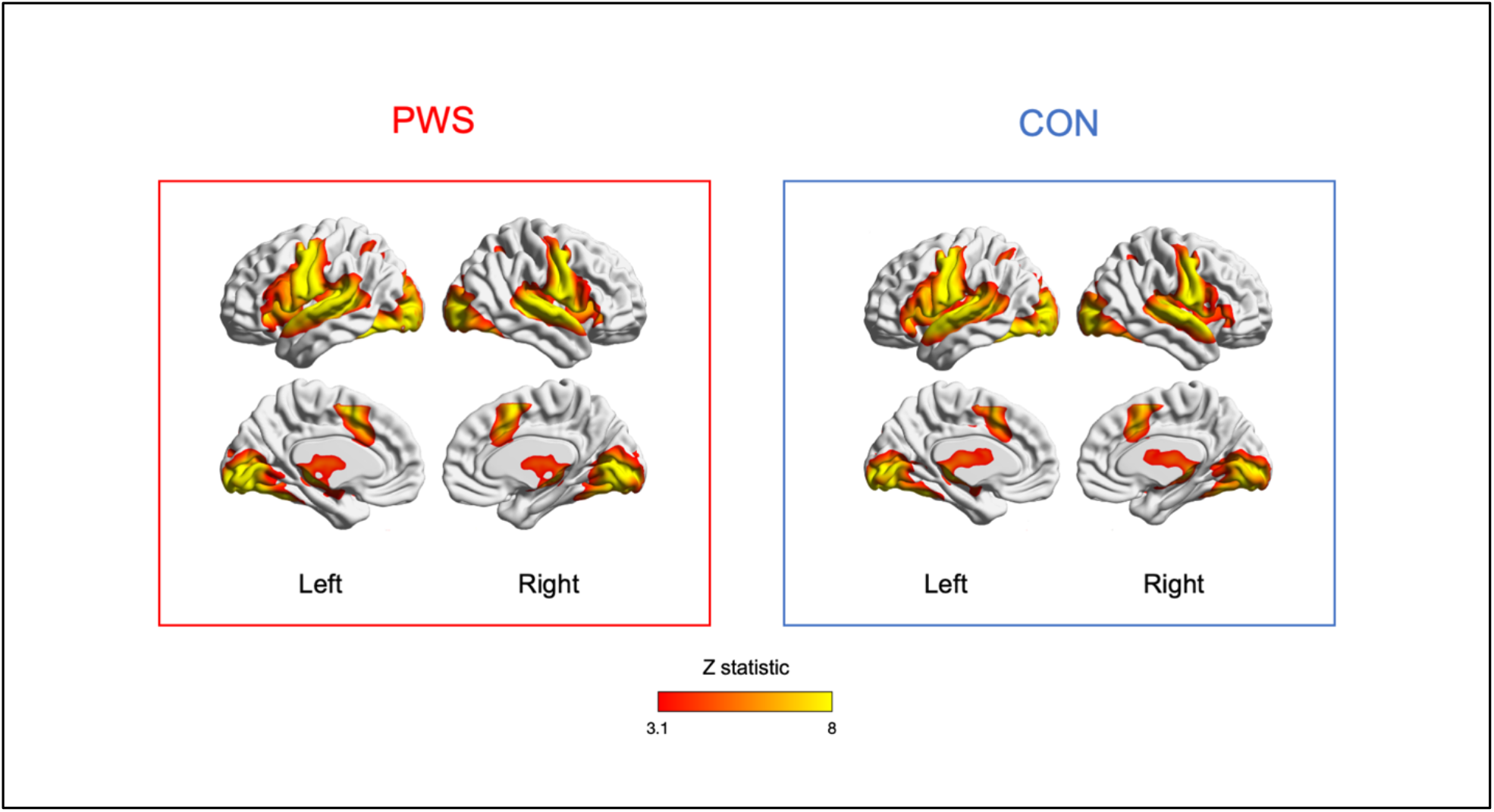
Brain activity during sentence reading. Mean BOLD activity during sentence reading compared with baseline rest conditions for the PWS and the control groups. Despite different speech fluency profiles, the patterns of brain activity were similar between the two groups. The data are represented on colour-coded statistical maps, which have been thresholded at a cluster-forming Z value greater than 3.1 and a cluster extent significance level of p < 0.05 (PWS = 56, CON = 53). These maps are superimposed on the MNI-152/Talairach surface template. The BrainNet Viewer (Xia et al., 2013) was used to visualise the medial and lateral surfaces of both the left and right cerebral hemispheres. The left column shows the PWS group, while the right column shows the control group. PWS: People who stutter; CON: Control group.

### Group differences

During the sentence reading task, no group differences survived whole-brain correction (using the cluster-forming threshold of Z > 3.1, and extent threshold p <0.05, corrected for family wise error). However, at an exploratory threshold (Z > 2.3, k > 30 voxels, uncorrected), differences in BOLD activity were observed between PWS and controls (Fig. 2; Table 2). PWS had greater activity compared with controls in the right frontal pole, right anterior insula extending to the frontal operculum, left planum temporale, and midbrain at the level of red nucleus bilaterally. Conversely, areas of decreased activity in PWS included the left superior frontal sulcus, subgenual medial prefrontal cortex, right anterior temporal lobe and portions of inferior parietal lobe bilaterally, including the left angular gyrus.

**Table 2:** Brain regions that exhibit differences in activity between people who stutter and controls during sentence reading. The differences are identified at an exploratory threshold with a cluster size exceeding 30 voxels. For each cluster the peak activation level, and the coordinates of the peak in MNI-152 standard space are provided. PWS = People who stutter; CON: Controls.

| Brain Region | X | Y | Z | # of Voxels | Z-statistic |
| --- | --- | --- | --- | --- | --- |
| *PWS greater than CON* |  |  |  |  |  |
| Right frontal pole | 32 | 52 | 24 | 96 | 2.87 |
| Right anterior insula | 38 | 20 | 6 | 101 | 3.25 |
| Left red nucleus | -6 | -20 | -10 | 35 | 2.81 |
| Right red nucleus | 10 | -22 | -14 | 62 | 2.93 |
| Left planum temporale | -54 | -36 | 12 | 62 | 2.85 |
| *CON greater than PWS* |  |  |  |  |  |
| Subgenual medial prefrontal cortex | -2 | 28 | -2 | 43 | 2.78 |
| Left superior frontal sulcus | -24 | 22 | 50 | 53 | 2.87 |
| Right anterior temporal lobe | 50 | 2 | -20 | 49 | 2.61 |
| Left angular gyrus | -44 | -56 | 32 | 74 | 3.09 |
| Right inferior parietal lobe | 58 | -66 | 20 | 120 | 3.17 |
| Left inferior parietal lobe | -36 | -70 | 28 | 131 | 2.75 |

**Figure 2:**
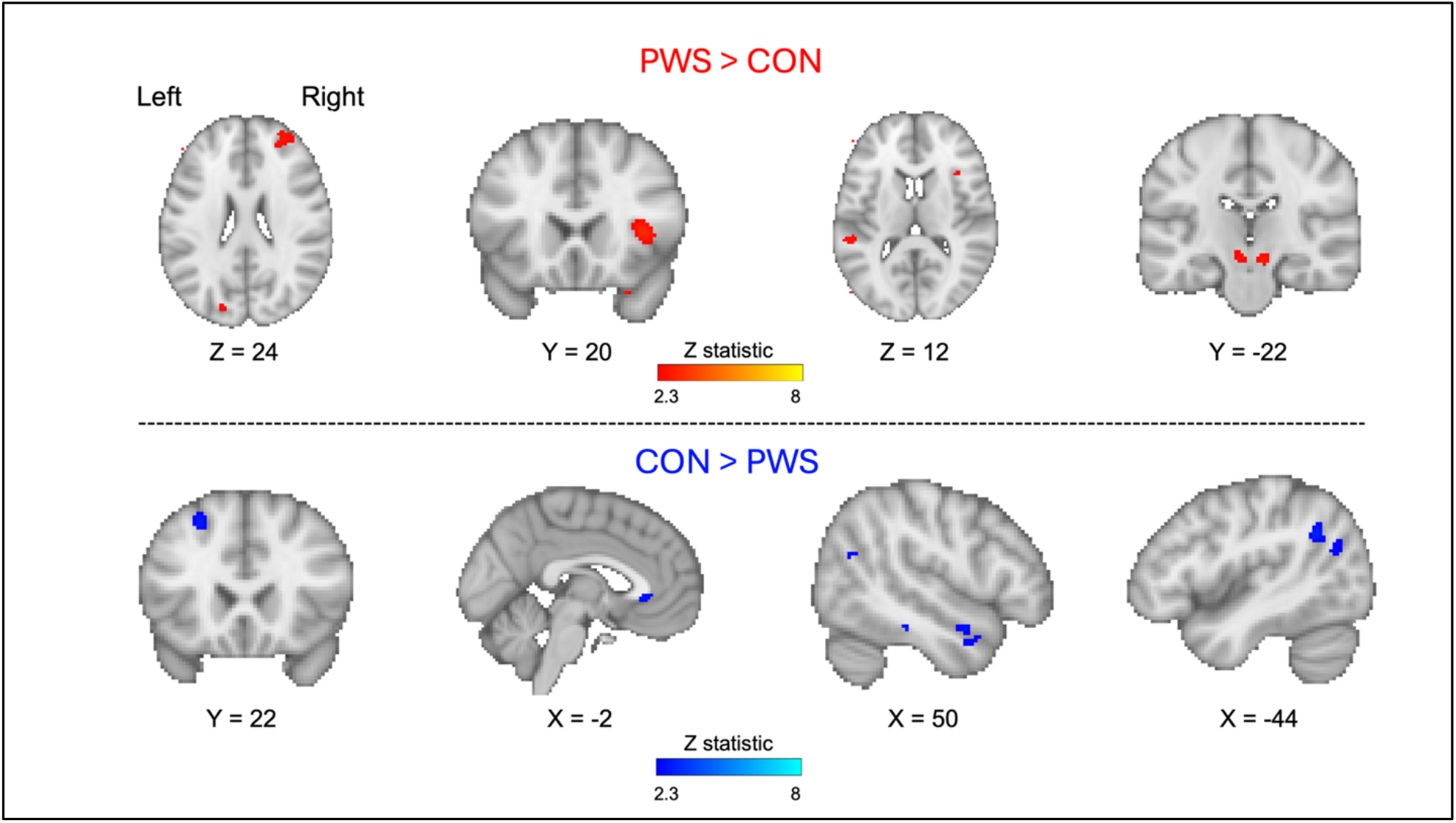
Differences in sentence-reading activity between people who stutter and controls. Coloured maps of the Z statistic are overlaid onto the MNI-152 average brain, with a threshold of Z > 2.3. Red regions indicate areas where PWS (n = 56) had significantly more activity than controls (n = 53) while blue regions indicate areas where PWS had less activity than controls (exploratory threshold of Z > 2.3, k > 30 voxels, uncorrected). For a detailed description of activity patterns, see main text. The numbers below each image indicate the coordinate in millimetres of that slice in x (for sagittal), y (for coronal), and z (for axial) relative to the orthogonal planes through the anterior commissure. PWS: People who stutter; CON: Control group.

## Discussion

We conducted an analysis combining four fMRI datasets with 109 participants, collected by our research group over the last two decades by using a consistent methodological approach and experimental task of sentence reading. This approach aimed to minimise the confounding effects often encountered in individual study designs within meta-analyses (e.g., different task, neuroimaging tool, methodology etc.), while leveraging a relatively large dataset to enhance the statistical power of our findings. Despite this larger sample size, our analysis did not reveal significant differences at a statistically robust level in BOLD activity in PWS compared with typically fluent controls. To avoid a potential false negative, we explored group differences at a lower uncorrected threshold. At this uncorrected threshold, we observed some findings that aligned with previous meta-analyses of functional imaging studies in stuttering. Of particular relevance, greater activity was detected in the right insular cortex in PWS, which was previously identified as one of the neural “signatures” of stuttering (Brown et al., 2005). However, no significant overactivity was observed in PWS in the cerebellum, nor was lower activity detected in superior posterior temporal lobes, both of which have also been suggested as neural markers (Brown et al., 2005). Subcortically, we found bilateral overactivity of the red nucleus in PWS, which contrasts with the findings of the meta-analysis by Budde and colleagues (2014), who reported reduced activity in the left red nucleus in PWS. The symmetrical pattern of overactivity of this nucleus in the two hemispheres is unlikely to be a false positive finding in our opinion.

Our analysis included stuttering events, mainly to increase the statistical power of our analysis. Furthermore, literature suggests that even during perceptibly fluent speech, PWS tend to exhibit greater variability in articulatory movements compared with controls (Wiltshire et al., 2021; Smith et al., 2010), complicating an accurate differentiation of stuttered versus fluent speech. Including stuttering events makes it more difficult to interpret the results, because previous research has mostly analysed only fluent speech. However, this approach likely gives a more representative depiction of the neural correlates of stuttering, especially since around 21 percent of sentence reading trials in our combined datasets included at least one stuttered utterance. The proportion of stuttered trials per participant ranged from 0% to 100%, and 92% of participants produced at least one stuttered trial.

Even though some of our datasets included additional conditions involving sentence reading using fluency enhancers such as metronome-paced speech and delayed auditory feedback, we excluded these data from our analysis, similar to the methodology employed by Budde and colleagues (2014) in their refined data analysis. The primary rationale for this exclusion was to maximise comparability across participants by retaining only the normal sentence reading condition with normal feedback, as activity patterns during fluency enhancement can differ significantly (Toyomura et al., 2011). These variations in activity patterns suggest the existence of multiple mechanisms for achieving fluency, which could potentially confound the interpretation of neural activity (Budde et al., 2014).

### Overactivity in the right anterior insula/frontal operculum

In the meta-analysis of Brown and colleagues (2005), overactivity in the right anterior insula/frontal operculum in PWS was identified as one of the neural signatures of stuttering. Subsequent meta-analysis by Budde and colleagues (2014) confirmed this finding by reporting increased activity in this region in PWS compared with controls. Furthermore, within-group analysis revealed elevated right anterior insula and frontal operculum activity specifically during stuttering events (Kell et al., 2009; Preibisch et al., 2003; Connally et al., 2018). An fMRI study contrasting imagined speech with imagined humming conditions in PWS pointed to increased activity in right hemispheric regions, such as the right posterior inferior frontal gyrus, specifically pars opercularis, which is part of the frontal operculum (Neef et al., 2018). Furthermore, the study found that the severity of stuttering was associated with the strength of white matter connections in these hyperactive right frontal regions. The authors interpreted this as stuttering severity being linked to enhanced structural connectivity of the motor response inhibition network.

Overall, overactivity in right anterior insula/frontal operculum could be interpreted in various ways: potentially causing disfluency through increased inhibition (Neef et al., 2018), manifesting errors in speech production (Tourville et al., 2008), or attempting to mitigate them (Kell et al., 2009). To further elucidate these hypotheses, the use of non-invasive brain stimulation methods such as ultrasound stimulation to inhibit neural activity in right anterior insula/frontal operculum while subsequently measuring speech fluency and brain activity, could shed light on the causal relationship.

To better understand how the right anterior insular cortex contributes to different stages of speech production over time, it is worthwhile reviewing studies that have used electrophysiological neuroimaging methods, as these provide higher temporal resolution compared with fMRI. A magnetoencephalography study revealed preparatory motor activity in the insular cortex, with right hemisphere dominance, approximately 160ms after visual cues (Kato et al., 2007). This occurred during a task where participants imagined articulating phonograms, suggesting that the right insula may be involved in early stages of speech preparation even without overt articulation. This aligns with findings from another study showing that the right insular cortex is involved in auditory verbal imagery (Pratts et al., 2023). It is important to note that the insular regions serve broader functions beyond speech. For example, the insula may adjust autonomic activity in response to respiratory demands during articulation, as supported by a PET study showing predominantly right-lateralised activation in the mid and anterior insula during dyspnea (shortness of breath) (Banzett et al., 2000). Our findings are consistent with the possibility that the elevated right anterior insula/frontal operculum activity reported here reflects differences in early articulatory planning, regulation of respiration during speech, or an interaction between the two. Direct evidence remains limited, and studies that combine laryngeal EMG or breathing measurements with high-resolution electrophysiology such as optically pump magnetometers (OPM) could help clarify these alternatives.

### Overactivity in red nucleus

The red nucleus contributes to motor control and evolutionary adaptations to varying motor demands across different species. In humans, our understanding of the connections of this nucleus mainly comes from region of interest-based resting-state fMRI analyses (Nioche et al., 2009; Zhang et al., 2015; Krimmel et al., 2025). A resting-state fMRI study described an extensive red nucleus functional connectivity network, including regions such as basal ganglia and insula (Zhang et al., 2015), which have been reported as key brain regions in stuttering (Watkins et al., 2008; Brown et al., 2005; Chang & Guenther, 2020; Theys et al., 2024). Extending these findings, a recent large-scale resting state fMRI study reported that the human red nucleus is functionally connected more with action-mode and salience networks than with primary motor regions, suggesting a role in integrating goal-directed action rather than direct motor execution (Krimmel et al., 2025). Further supporting its role in volitional motor control, task fMRI studies with healthy participants have reported stronger activity in the red nucleus during the preparation of self-initiated finger movements compared with externally triggered movements (Boecker et al., 2008; Cunnington et al., 2002). For instance, one study reported increased activity during the planning phase of the internally initiated finger movement condition, as opposed to the externally triggered condition (Boecker et al., 2008). This increased activity was also observed in the frontal regions, including dorsolateral prefrontal cortex, inferior parietal regions, and insula, which align with the differences in activity we observed in PWS during our self-initiated sentence reading task. Therefore, one possible interpretation could be that the increased activity in the red nucleus during the planning phase of self-initiated motor movements might represent the heightened effort required to execute speech motor commands during sentence reading in PWS.

In chronic post-stroke patients, strengthening of the cortico-rubro-spinal network, including the red nucleus, may serve as a compensatory mechanism for pyramidal tract damage, and aid motor recovery. This hypothesis was supported by reported correlations between fractional anisotropy values (a metric for white matter integrity), and motor function assessments, suggesting that enhanced integrity in this pathway correlates with improved motor performance (Rüber et al., 2012). Specifically, increased fractional anisotropy in cortico-red nucleus tracts have been positively correlated with the level of motor impairment in chronic post-stroke patients at various intervals post-lesion, indicating plasticity of this region may compensate for the affected pyramidal tracts (Jang & Kwon, 2015; Kim et al., 2018; Rüber et al., 2012) and taking over part of the motor command normally conveyed by them and thus compensating for their loss. A similar compensatory logic might apply to stuttering, as it has been linked to structural and functional anomalies in the cortico-basal ganglia-thalamocortical loop (Watkins et al., 2008; Giraud et al., 2008; Wu et al., 1997; Theys et al., 2024), that supports self-initiated movement (Alm, 2004; Burghaus et al., 2006; Chang & Guenther, 2020). When this primary circuit is inefficient, increased red-nucleus activity might provide a compensatory subcortical route. Consistent with this proposition, our findings revealed increased activity of the red nucleus bilaterally, extending the results of Budde and colleagues (2014), who reported lower activity in the left red nucleus, derived exclusively from epochs of fluent speech. Nevertheless, the precise contribution of the red nucleus to stuttering remains to be defined, but converging evidence supports its dynamic involvement.

## Limitations and Future Directions

Our study has several limitations that must be considered when interpreting the findings. Firstly, we observed sub-threshold differences in activity in brain regions that are critical for self-initiated motor commands, such as the anterior insula/frontal operculum, and the red nucleus. However, interpreting these results is challenging due to the inherently low temporal resolution of fMRI. This limitation restricted our ability to precisely map the dynamic temporal processes involved in stuttering.

Secondly, the ecological validity of using a sentence reading task to study stuttering is limited. Stuttering is more likely to occur during spontaneous information transfer when there is communicative intent, rather than in isolated reading tasks (Jackson et al., 2021). Future studies may aim to enhance ecological validity by incorporating artificial intelligence-driven conversational agents during scanning, or real-time online conversations with people during the scan, to prevent isolation in the scanner. This approach would facilitate real-time interactions, more closely mirroring everyday communication and potentially providing deeper insights into the neural correlates of stuttering.

Finally, it is worth mentioning the potential for methodological confounds that may bias findings towards false negatives. Variability in neural activity patterns among PWS, such as differences in fluency or compensatory strategies, may mask region-specific effects at the group level. Future studies could address this by recruiting people with more severe stuttering to enhance the heterogeneity of the sample. Additionally, the sparse sampling design, although effective at reducing noise and capturing the haemodynamic response peak, possibly limits temporal resolution. Moreover, combining data from studies that used different acquisition parameters for spatial resolution introduces additional measurement variance, which may further mask subtle group effects.

## Conclusions

In this combined analysis of four fMRI studies, we investigated the neural correlates of stuttering using a relatively large sample. The studies included a consistent methodological approach and task of sentence reading. Although group differences in task-evoked activity did not remain significant after correction for multiple comparisons, we observed some results at a lower uncorrected threshold that were consistent with earlier meta-analyses of functional imaging studies in stuttering. For example, we found increased activity in right anterior insular cortex, reaffirming its role as a neural marker of stuttering, and bilateral overactivity in the red nucleus of PWS, which may reflect either a compensatory mechanism for disruptions in the speech motor pathway or a disruptive influence on fluency. These findings highlight the complex neurobiological mechanisms underlying stuttering, involving both cortical and subcortical regions. While our study advances the understanding of the neural correlates of stuttering, it also emphasises the need for further research to explore these differences causally and to inform the development of therapeutic interventions.

## Abbreviations

PWS: people who stutter;
TFS: typically fluent speakers;
LI: laterality index;
ROI: region of interest.

## Acknowledgements

We are deeply grateful to the Dominic Barker Trust for their support of our research. We also extend our heartfelt thanks to all the participants who took part in this study. To ensure open access, the author has applied a CC BY public copyright license to any Author Accepted Manuscript version resulting from this submission.

## Data Availability

The brain imaging data of the group averages in the standard space are available at NeuroVault: https://neurovault.org/collections/21188/

